# Derivation of a Time-Dependent Model for Long-Term Cortical Bone Adaptation

**DOI:** 10.1101/2025.08.31.673332

**Authors:** Jitendra Prasad

**Affiliations:** Department of Mechanical Engineering, Indian Institute of Technology Ropar, Rupnagar, Punjab, India - 140001

## Abstract

This work attempts to derive long-term, time-dependent cortical bone adaptation to mechanical loading. Linear control theory is used to model the adaptation process, with the stimulus defined in terms of dissipation energy density. The newly adapted area is expressed as a function of the stimulus in the form of a differential equation, which is analyzed to obtain closed-form solutions. The study explores different possibilities, such as varying the order of differential equations (including fractional order) and examining different types of responses, e.g., critically damped and overdamped. Such model diversity will help identify the most appropriate formulation that fits experimental data.

## 1. Introduction

Bone is a living tissue that continuously adapts to its mechanical environment via remodeling mechanisms driven by osteocytes, osteoblasts, and osteoclasts – ensuring structural integrity and metabolic balance. The foundational concept of Wolff’s law posits that bone architecture adapts according to mechanical loading [1]. Modern understanding emphasizes that osteocytes act as mechanosensors translating mechanical stimuli into biochemical signals that regulate bone formation and resorption [2], [3].

Various mathematical models have been developed to capture bone adaptation. Early phenomenological models based on strain or strain energy density provided useful insights [4], [5], [6], [7]. With advances in biomechanics and computational methods, formulations that include poroelastic behavior, viscoelasticity, and finite element simulations have emerged to better characterize bone’s hierarchical response [8], [9], [10], [11], [12], [13]. However, these models often fall short in predicting long-term adaptation under sustained or repeated mechanical loading.

To address that gap, control theory has emerged as a promising modeling framework for bone mechanobiology [14], [15]. This approach allows physiological processes – such as ATP release in osteocytes, calcium signaling, and collagen deposition – to be represented as interconnected subsystems, leading to higher-order differential equations describing cross-sectional bone adaptation. Moreover, using dissipation energy density (DED) as the stimulus provides a physically grounded measure linking mechanical loading with osteocytic transduction [9], [10], [16], [17].

Building on this, Shekhar et al. recently demonstrated that modeling cortical bone resorption in a mouse tibia under disuse conditions using poroelastic finite element analysis and dissipation energy density can accurately predict site-specific bone loss [10]. Their model revealed that the mineral resorption rate is proportional to the square root of the loss of DED [10].

Additionally, Singh et al. presented a novel mathematical derivation and validation of loading-induced new bone formation using mineral apposition rate (MAR) as the response variable, reinforcing the value of mechanistic modeling approaches using dissipation energy density [9]. The current study extends that foundation by modeling long-term cortical bone adaptation under repeated loading, incorporating overdamped, critically damped, and fractional (rational) order formulations to better match experimental dynamics.

In this paper, linear control theory with dissipation energy density as the stimulus is employed to derive models of bone cross-sectional area adaptation under mechanical loading, exploring both integer- and rational-order systems. This diverse modeling framework aims to closely match experimental observations and provide deeper insights into the temporal dynamics of cortical bone adaptation.

## 2. Methods

### 2.1. Dissipation Energy Density

Let us assume that the bone is subjected to a normal stress *σ*(*t*), which produces a normal strain *ε*(*t*).

The total dissipation energy density for time *T*is given by [9], [16]:

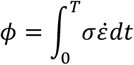

For example, for a sinusoidal stress:

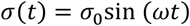

and the corresponding strain:

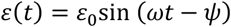

Where

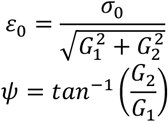

and *G*_1_ + *iG*_2_ is the complex modulus of the bone.

For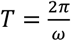, the dissipation energy density is:

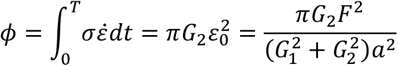

where *F* is the normal force and *a* is the area of cross section.

For the same force or loading protocol:

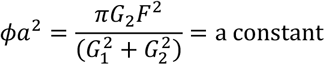

or equivalently:

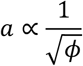

which leads to:

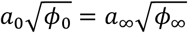

where the subscripts 0 and ∞ correspond to time *t* = 0 and *t* = ∞, respectively.

### 2.2. Conservation of the Time-Average of Dissipation Energy Density

#### Statement

If a loading protocol is initiated and maintained indefinitely, the bone cross-section will adapt such that the final (steady state) area of cross-section of the bone (*A*_∞_) ensures that the average dissipation energy density is maintained at a universal value *ϕ*_∞_.

In other words,

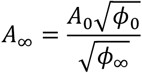

### 2.3. Derivation

#### 2.3.1. Control Theory Approach to Bone Formation Process

Bone adaptation involves many processes. For example, mechanical loading leads to energy dissipation, which results in APT release in osteocytes, triggering calcium ion signalling that reaches osteoblasts. Eventually collagen is deposited and mineralized by the osteoblasts.

Each process (say, the *i*-th process) has an input function *r*_*i*_(*t*) and an output *c*_*i*_(*t*) . In linear control theory, each process has a transfer function, defined as the ratio of the output *C*_*i*_(*s*) to the input *R*_*i*_(*s*), in the Laplace domain. For example, a first-order system with delay time relates output *c*_*i*_(*t*) and input *r*_*i*_(*t*) through a transfer function *F*_*i*_(*s*) as follows:

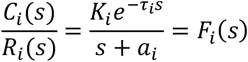

The corresponding differential equation relating input and output is:

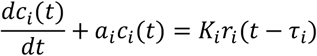

If there are *n* such processes in series in new bone formation – where output of the preceding process (the (*i*-1)-th) acts as the input of the current process (the *i*-th), and the output of the current process acts as the input of the following process (the (*i*+1)-th) – then the following relationship holds:

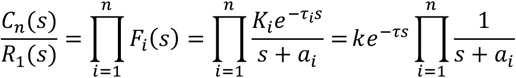

The corresponding *n*-th order differential equation is:

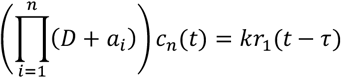

In bone adaptation, the output *c*_*n*_(*t*) may be taken as the area of the new bone formed, *A*(*t*), at a cross-section of interest, and the input *r*_1_(*t*) may be taken as the mechanical stimulus, *f*(*t*), applied to the osteocytes. The revised equation becomes:

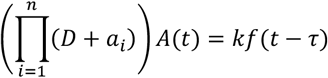

Here, the time *t* is macroscopic (e.g., measured in weeks), as bone takes weeks to adapt. If a particular weekly loading protocol is maintained indefinitely, i.e. the same stimulus (*f*_0_) is repeated every week, then the stimulus *f*(*t*) is given by:

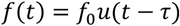

where *u*(*t*) is the unit-step function.

The final form of the differential equation is:

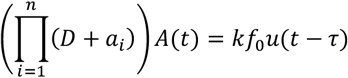

We will analyse two cases:

i. *a*_*i*_’s are distinct, i.e., the system is overdamped, and
ii. All *a*_*i*_’s are the same (*a*_*i*_ = *α*), i.e., the system is critically damped.

#### 2.3.2. The n-th Order Overdamped System

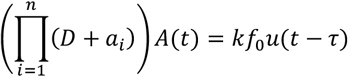

where

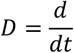

Taking Laplace transform of both sides:

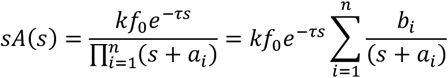

Thus,

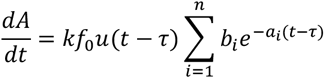

or equivalently,

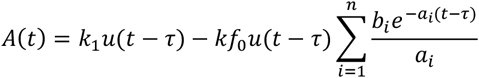

Applying initial and end conditions:

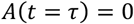

And

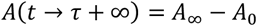

At *t* = *τ*,

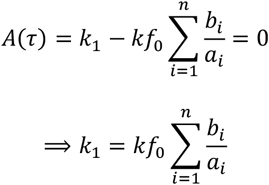

Hence,

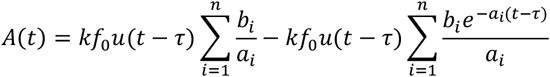

Also,

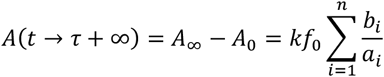

or

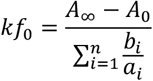

Therefore,

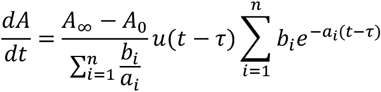

However, using the conservation relation:

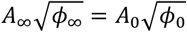

we obtain:

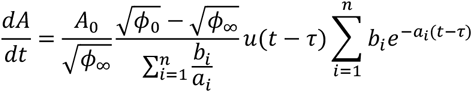

or equivalently,

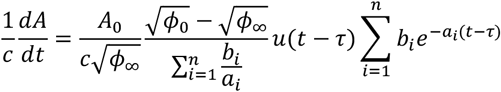

where *c* is the circumference of the bone cross-section.

Defining the bone formation rate (BFR):

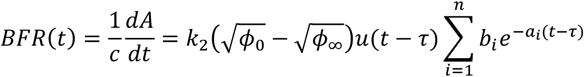

where

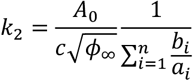

and *k*_2_ is a constant.

This formulation shows that the bone formation rate (BFR) is proportional to the difference between the initial and final (universal) dissipation energy densities 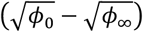. The BFR depends on the exponential decay terms associated with each sub-process of bone adaptation. Initially, BFR is dominated by the transient response of the system, whereas at long times it asymptotically approaches a steady state value of zero.

#### 2.3.3. n-th Order Critically Damped System

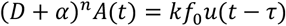

Applying the Laplace transform to both sides, we get:

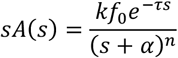

On inverse Laplace transform:

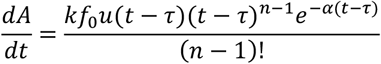

Subject to the conditions:

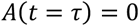

and

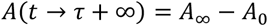

The above equation may be solved to obtain *A*(*t*).

Alternatively,

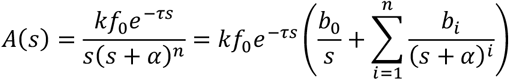

or equivalently,

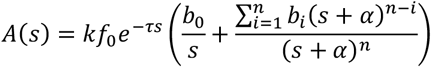

where:

i. The coefficient of *s*^*n*^:

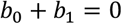
ii. The constant term:

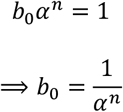

On inverse Laplace transform:

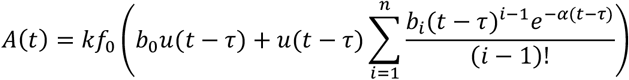

At steady state:

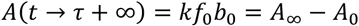

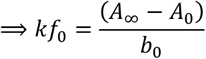

Therefore,

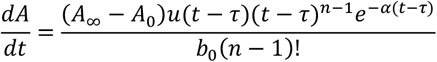

Using the relation

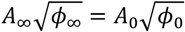

we obtain:

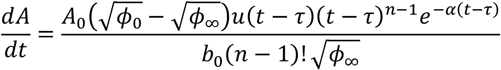

or equivalently, defining the bone formation rate (BFR):

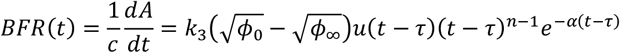

where

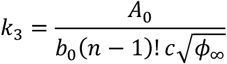

or equivalently,

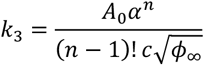

and *k*_3_ is a constant.

In the critically damped case, the bone formation rate (BFR) exhibits a smooth, non-oscillatory transient response. It initially rises after the delay *τ*, peaks due to the polynomial term (*t* − *τ*)^*n*−1^, and then decays rapidly under the exponential term *e*^−*α*(*t*−*τ*)^. Over time, BFR approaches zero, indicating that the system stabilizes at the new adapted bone cross-sectional area without overshooting or oscillations.

#### 2.3.4. Fractional / Rational Order Critically Damped System

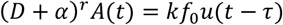

where *r* = *n* + *β* and 0 < *β* < 1.

Applying the Laplace transform:

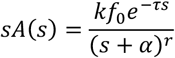

On inverse Laplace transform:

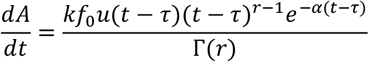

where Γ(*r*) is the Gamma function.

Initial and final conditions,

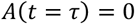

and

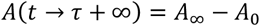

The new bone formation area *A*(*t*) can also be obtained from the inverse transform of:

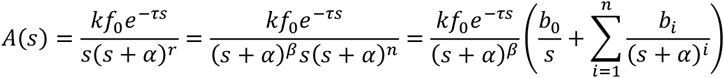

where, as in the previous section,

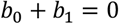

And

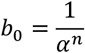

Rearranging:

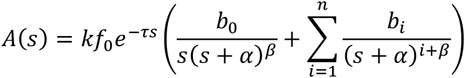

By inversing Laplace transform:

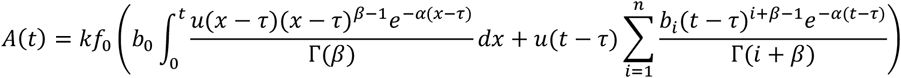

At steady state:

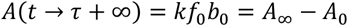

which gives:

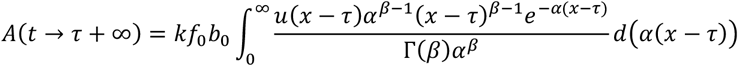

Or equivalently,

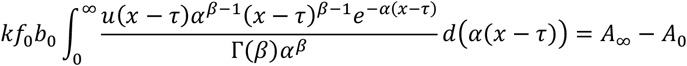

and finally:

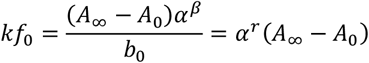

Therefore,

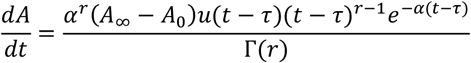

or equivalently, defining the bone formation rate (BFR), as earlier:

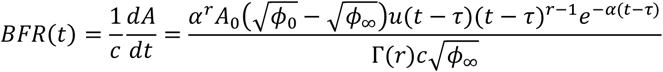

Thus,

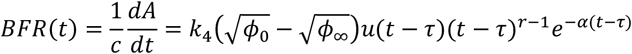

where

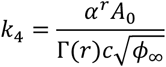

and *k*_4_ is a constant.

The fractional (or rational) order formulation generalizes the critically damped system by allowing the order of the differential operator to take non-integer values. This introduces intermediate dynamics between two consecutive integer-order critically damped responses. In the context of bone adaptation, the fractional order model can capture more realistic physiological processes where adaptation is neither purely exponential nor purely polynomial, but instead exhibits non-integer power-law–like behavior characteristic of complex biological systems.

## 3. Results and Discussion

### 3.1. First-Order System

Assuming bone formation as a first order system, the expression for the BFR reduces to:

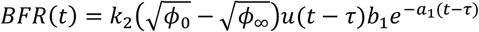

This is plotted in Fig. 1, assuming 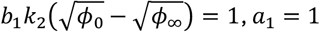 and *τ* = 1 week.

**Figure 1.**
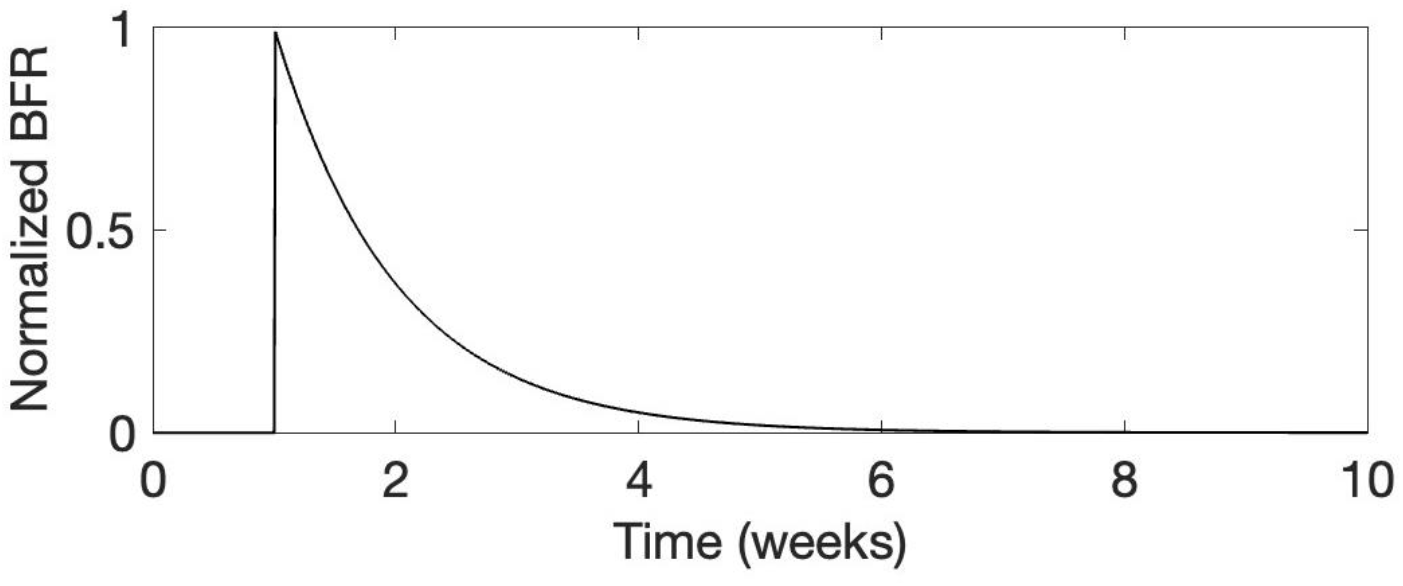
Normalized bone formation rate (BFR) assuming a first-order system

### 3.2. Second-Order Overdamped System

The bone formation rate (BFR) corresponding to the second-order system reduces to the following expression:

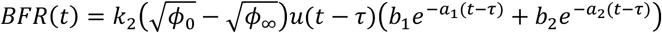

Imposing the condition *BFR*(*τ*) = 0 :

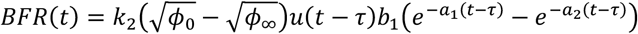

Assuming 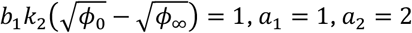, and *τ* = 1 week, the BFR is plotted in Fig. 2.

**Figure 2.**
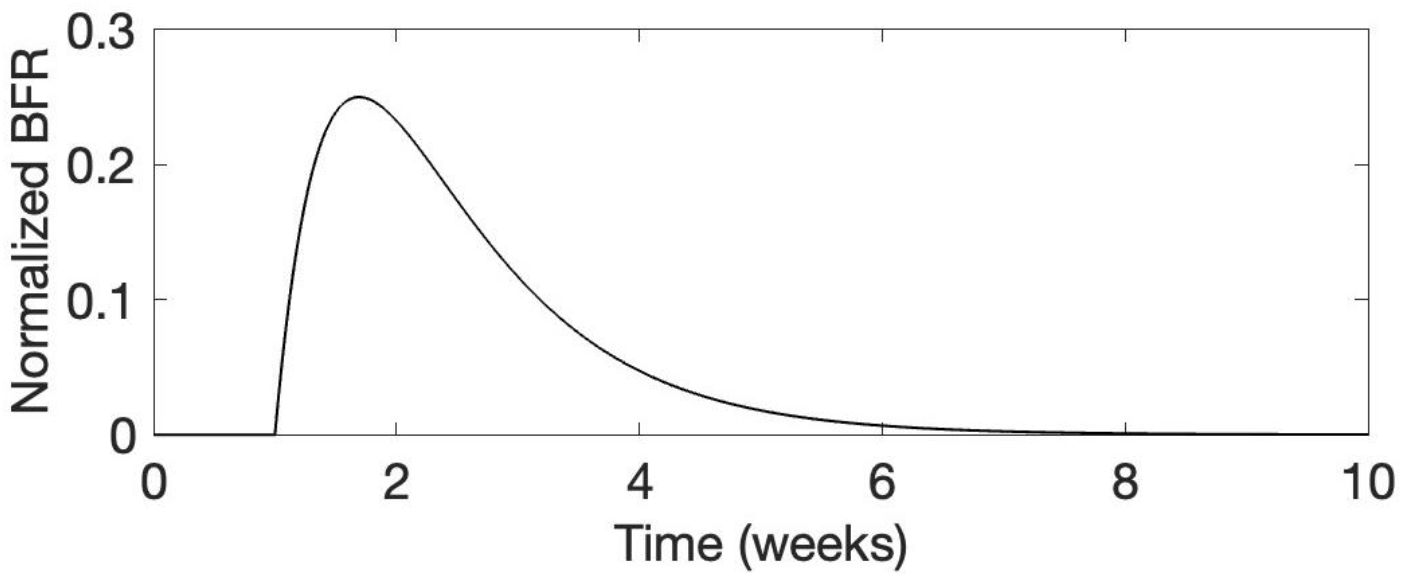
Representative bone formation rate assuming a second order system

### 3.3. Second-Order Critically Damped System

For the second-order critically damped system, the BFR is given by:

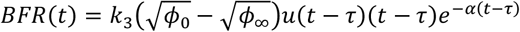

For 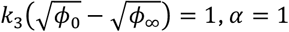 and *τ* = 1 week, the BFR is plotted in Fig. 3.

**Figure 3.**
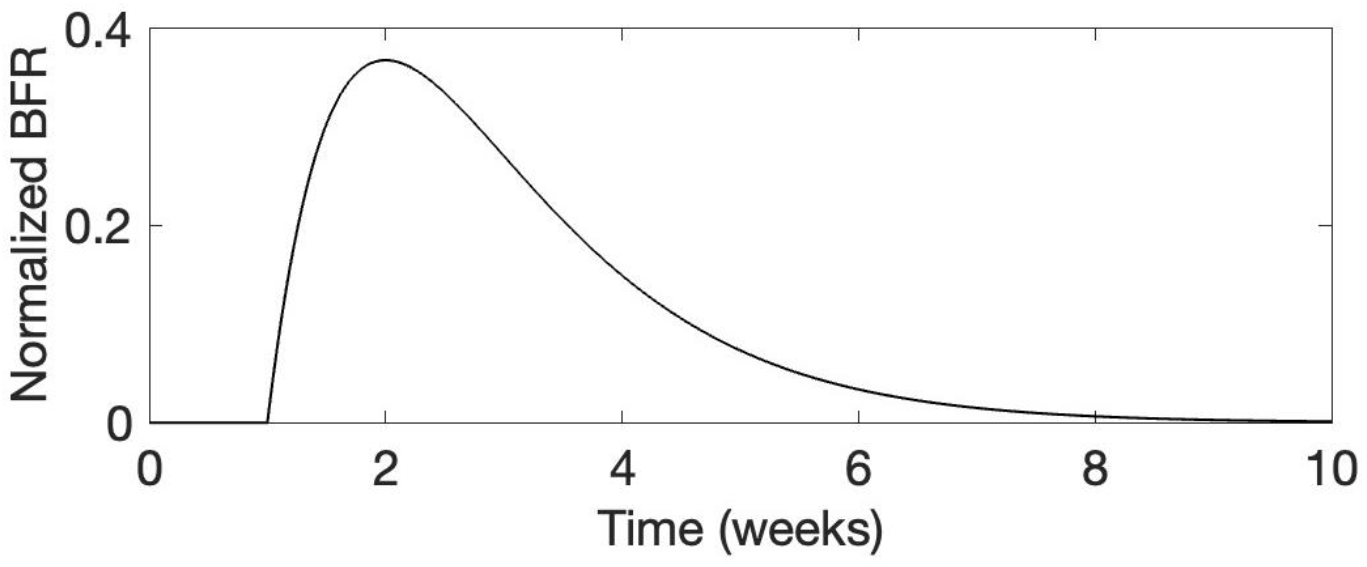
BFR assuming a second-order critically-damped system

### 3.4. Third-Order Critically Damped System

For the third-order critically damped system, the BFR is:

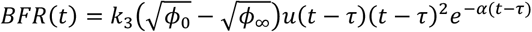

The corresponding plot (for the same parameters as in the previous section) is shown in Fig. 4.

**Figure 4.**
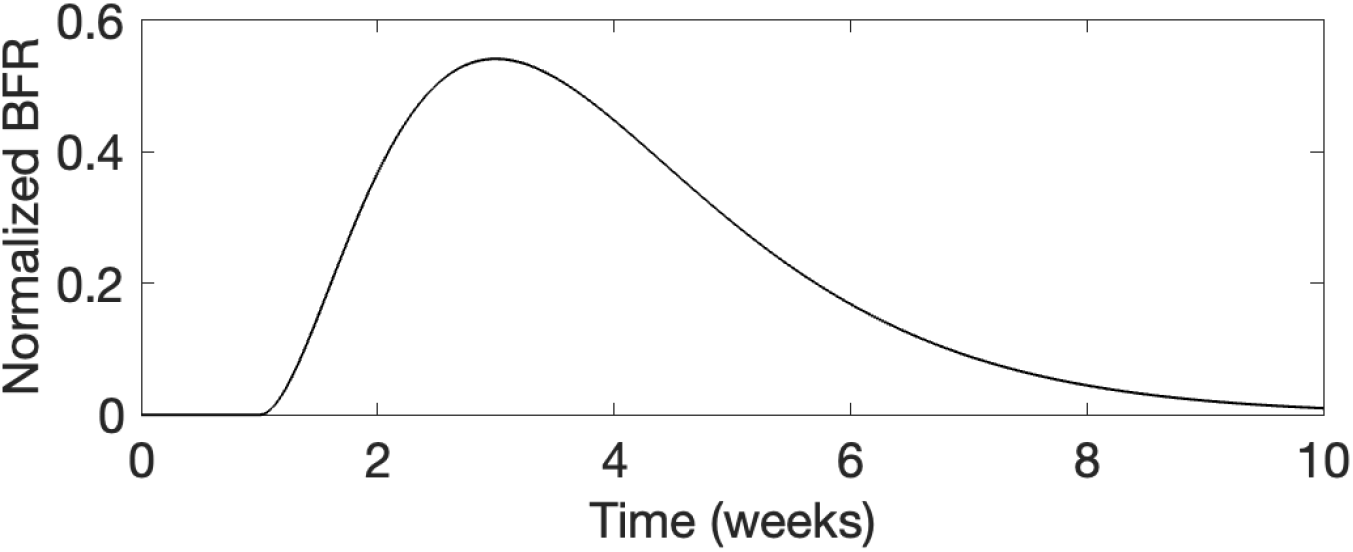
BFR for a third-order critically damped system

### 3.5. Two-And-a-Half Order Critically Damped System

The 2.5-order critically damped system corresponds to a fractional (rational) order system with *r* = 2.5. The corresponding BFR is:

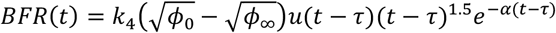

The BFR is plotted in Fig. 5 for the following parameters: 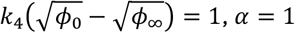 and *τ* = 1 week.

**Figure 5.**
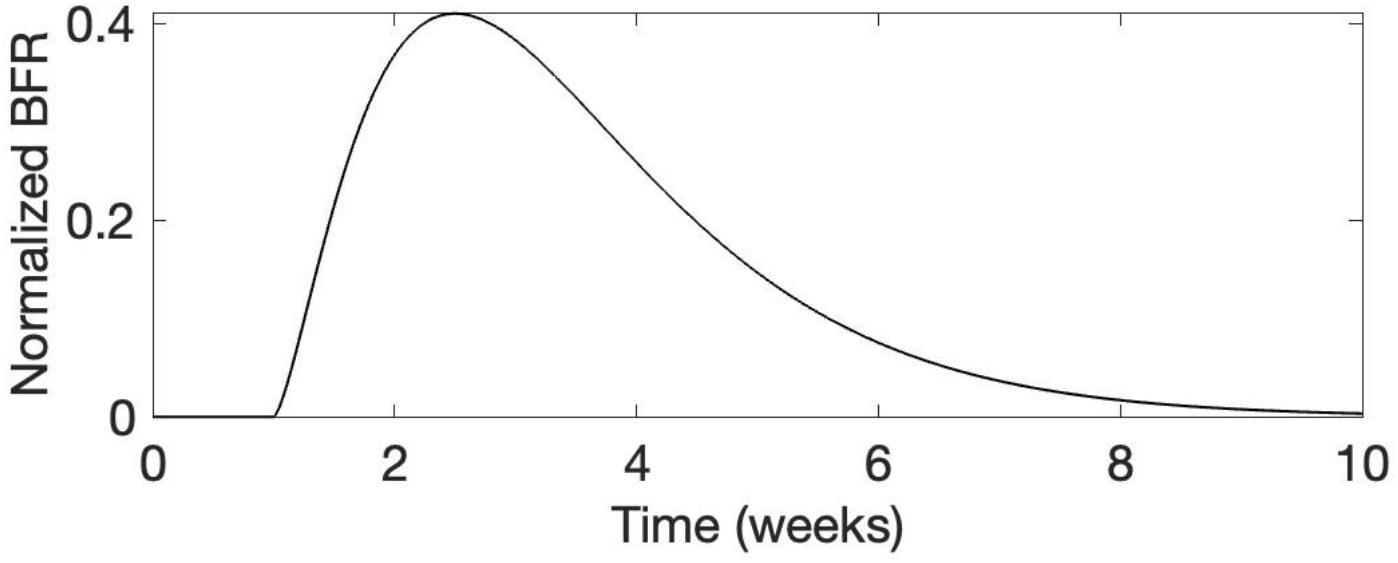
BFR for the 2.5-order system

### 3.6. Three-And-a-Half Order Critically-Damped System

If the bone-adaptation system is assumed to be of order of 3.5, the corresponding equation for BFR is:

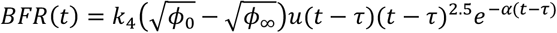

The BFR for this system is shown in Fig. 6.

**Figure 6.**
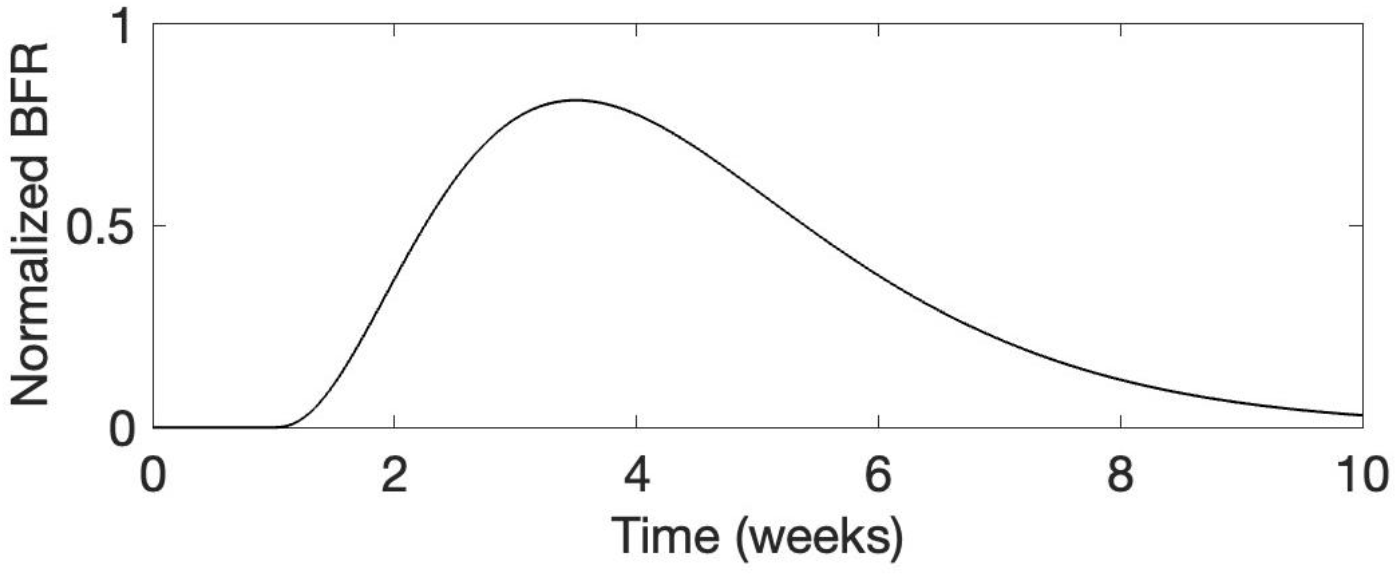
BFR for the 3.5-order system

### 3.7. Comparison and Discussion

BFRs for all the cases discussed in the earlier subsections are compared in Fig. 7.

**Figure 7.**
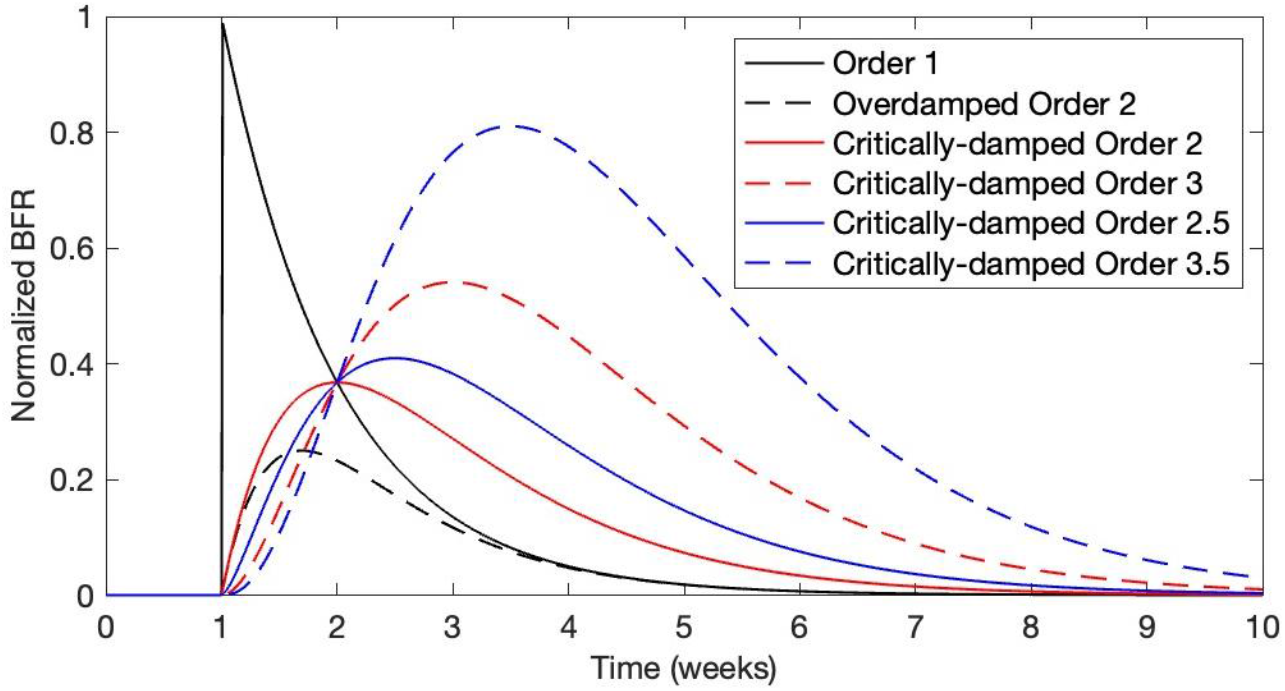
Comparison of all cases studied

From the comparison, it is observed that the first-order system is different from all others because BFR cannot be zero at *t* = *τ*. While BFR value for a critically damped system is always zero at *t* = *τ*, the overdamped system has the flexibility to make the BFR either zero or non-zero. However, experimental studies have confirmed that BFR slowly increases from zero to a peak and then gradually decreases towards zero [18]. Hence, the first-order system is not appropriate to model the bone adaptation process.

In all cases, one common feature is that BFR is directly proportional to 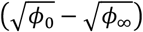. In fact, this proportionality was already derived in the author’s previous work [9], [16], assuming a first-order system. For a long-term, time-dependent system, however, the assumption of a higher-order system is more appropriate. The exact order needs to be determined by fitting experimental observations.

It can be seen that the BFR expression for integer-order critically-damped system is similar to that for the fractional (rational) order system, except for the appearance of (*n* − 1)! and Γ(*r*) in the two cases, respectively. This is expected, since Γ(*n*) = (*n* − 1)! In other words, the fractional (rational) order system converges to an integer-order system when *r* is an integer.

The applicability of higher-order models (including whether overdamped or critically damped) needs to be further tested against experimental studies.

### 3.8. Extension of Results

The expression for BFR corresponds to the step-function stimulus as input, i.e., the stimulus is maintained on a weekly timescale. This result can be extended to two cases: (i) only one week of loading, and (ii) varying weekly stimulus.

#### 3.8.1. Only One Week of Loading

When there is only one week of loading, the input becomes *k f* _0_ *δ* (*τ*) in place of *kf*_0_*u*(*t* −*τ*), where *δ*(*τ*) is the Dirac-delta function, related to the unit step function *u*(*τ*) as:

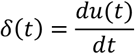

Since the governing differential equation is linear, the BFR corresponding to *kf*_0_*δ*(*τ*) (referred to as *BFR*_*δ*_(*t*) henceforth) will simply be the time derivate of the BFR corresponding to *kf*_0_*u*(*t* − *τ*) (referred to as *BFR*_*u*_(*t*) hereafter). For example, for the n-order overdamped system, the new BFR is:

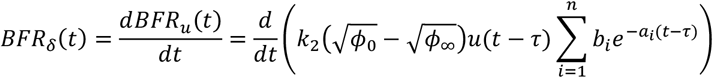

Similarly, the new BFR can be derived for other cases.

#### 3.8.2. Varying Weekly Stimulus

Once *BFR*_*δ*_(*t*) is known, it can be extended to determine the BFR corresponding to a varying stimulus (*BFR*_*x*_(*t*)) by convolution:

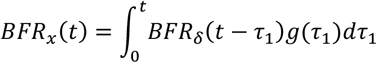

Where

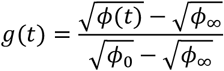

Here, *ϕ*(*t*) is the instantaneous dissipation energy density calculated with respect to the original cross-sectional area *A*_0_ , and *ϕ*_0_ is the dissipation energy density at the first week (i.e., at *t* = 0).

## 4. Conclusion and Future Work

The present work derived the bone formation rate (BFR) based on control theory. Three cases were considered: the integer-order overdamped system, the integer-order critically damped system, and the fractional/rational-order critically damped system. Together, these cover a wide spectrum of the time evolution of BFR, providing many parameters that can be tuned to fit experimental data in the future. The derived BFR expressions can not only model sustained stimulus but can also be extended to determine BFR for brief loading applications or even for varying loads on macroscopic timescale.

